# Temperature-induced compensatory growth in *C. elegans* is regulated by a thermosensitive TRP channel and influences reproductive rate

**DOI:** 10.1101/2020.12.08.415851

**Authors:** Zuzana Sekajova, Elena Rosa, Foteini Spagopoulou, Panagiotis-Ioannis Zervakis, Martin I. Lind

## Abstract

1. Animals are often not growing at the maximum rate, but can compensate for a bad start of life by subsequently increasing growth rate. While this compensatory growth is widespread, its direct fitness consequences are seldom investigated and its genetic basis is unknown.
2. We investigated the genetic regulation, as well as fitness and lifespan consequences of compensatory growth in response to temperature, using *C. elegans* knockout of the thermo-sensitive TRP ion channel TRPA-1, involved in temperature recognition. We exposed juvenile worms to cold, favourable (intermediate) or warm temperatures in order to delay or speed up development.
3. Wild-type worms initially exposed to cold temperature experienced slower growth but after being switched to a more favourable temperature, they expressed compensatory growth and caught up in size. Those initially reared at warmer temperatures than favourable experienced slower growth and attained smaller adult size after being switched to the most favourable temperature.
4. Compensatory growth also altered the reproductive schedule. While rate-sensitive individual fitness decreased by cold juvenile temperatures, as a direct effect of the substantial developmental delay, once worms returned to more favourable temperature, they shifted their reproductive schedule towards early reproduction. Therefore, when focusing on the post-treatment period, the reproductive rate increased even though lifetime reproductive success was unaffected. Surprisingly, compensatory growth did not reduce adult lifespan. In contrast to the findings for wild-type worms, juvenile temperature did not induce compensatory or slowed-down growth in the *trpa-1* knockout mutants.
5. We thus show that the *trpa-1* is involved in the network regulating temperature-induced compensatory growth in *C. elegans* and that this compensatory growth can influence the reproductive rate.

## Introduction

Adaptive phenotypic plasticity is the ability of a genotype to alter its phenotype depending upon the environmental conditions, in order to increase performance in a specific environment (Dewitt & Scheiner, 2004). For example, a common observation is that organisms faced with poor juvenile conditions (such as low temperature or food availability) are constrained to a low early-life growth rate, which can be increased in later life stages, if conditions improve, as a plastic response to compensate for the bad start. Importantly, such poor-condition individuals can have a higher growth rate than individuals that were not restricted as juveniles, which is termed compensatory growth (Hector & Nakagawa, 2012; Jobling, 2010). Compensatory growth often results in catch-up growth which is attainment of the same size compared to the non-restricted individuals (however, catch-up in size is not always reached by compensatory-growth, for example if the same size is reached solely by a longer development) (Hector & Nakagawa, 2012). Examples of compensatory growth are widespread (Hector & Nakagawa, 2012); it is present in organisms such as mice (Ozanne & Hales, 2004), frogs such as common frog (*Rana temporaria)* (Burraco et al., 2020; Orizaola et al., 2014) and moor frog (*Rana arvalis)* (Murillo□Rincón et al., 2017), zebra finches *(Taeniopygia guttata)* (Fisher et al., 2006) and fish such as sticklebacks *(Gasterosteus aculeatus* and *Pungitius pungitius* respectively) (Ab Ghani & Merilä, 2015; Lee et al., 2013), green swordtails (X*iphophorus helleri*) (Royle et al., 2006) and salmon (*Salmo salar*) (Morgan & Metcalfe, 2001). Typically, compensatory growth is expected to be adaptive in environments with a limited window for development and reproduction, such as time-constrained environments with strong seasonality, where a delayed development can be very costly (Metcalfe & Monaghan, 2003; Orizaola et al., 2014). In contrast to compensatory growth, its opposite, i.e. slowed-down growth or negative compensation, can happen after a return to standard (i.e. within the range usually experienced by animals) conditions after a period of environmentally induced increase in juvenile growth (Lee et al., 2013). This also suggests that there is a cost to the fast growth, therefore organisms which appear ahead may benefit from slowing growth rate and thus limiting the cost, once the environment returns to normal (Lee et al., 2013). In contrast to compensatory growth, very little attention has been given to slowed-down growth.

Despite an extensive research interest in compensatory growth plasticity, studies mostly focus on growth and associated costs in relation to lifespan (Metcalfe & Monaghan, 2003) and recently adult physiology (Burraco et al., 2020; Murillo□Rincón et al., 2017). However, reproductive performance is an essential element for understanding of the life-history consequences of compensatory growth, but is seldom investigated (notable exceptions are Auer et al., 2010; Dmitriew & Rowe, 2011; Lee et al., 2012). Moreover, while we now understand the selective pressures operating on the evolution of compensatory growth plasticity and some costs have been identified, its genetic regulation remains unknown.

Studies in model organisms have identified the evolutionarily conserved nutrient-sensing pathways (i.e. insulin/insulin-like growth factor signaling [IIS] and mechanistic target-of-rapamycin [mTOR]) as key regulators of life-history traits such as growth, reproduction and lifespan across animals (Dantzer & Swanson, 2012; Durmaz et al., 2019; Flatt & Heyland, 2011; Fontana et al., 2010; Lind et al., 2017; Regan et al., 2020). Importantly, these nutrient-sensing pathways can also underlie adaptive phenotypic plasticity. While IIS/mTOR are well known to respond to nutrient status, they are also key life-history regulators that take input from a range of environmental cues, including photoperiod and temperature (Feeney et al., 2016; Metaxakis et al., 2014; Sim et al., 2015). For example, insulin signaling in response to temperature regulates body size plasticity in *Hydra* (Mortzfeld et al., 2019), and is involved in body size polyphenism in horned beetles (Snell-Rood & Moczek, 2012) and in ovariole number in *Drosophila* (Green & Extavour, 2014). Therefore, it has recently been proposed that IIS/mTOR have evolved as general and conserved regulators of adaptive phenotypic plasticity (Regan et al., 2020), since they integrate upstream environmental cue recognition and downstream signaling into a physiological response. Therefore, these pathways and genes could also be part of the proximate mechanisms underlying adaptive compensatory growth.

Two common environmental factors that result in a poor start in life are low food availability (Hector & Nakagawa, 2012) and/or in case of ectotherms, low temperature (Lee et al., 2013). The latter can be less invasive to study experimentally, as it does not alter the nutritional intake per se (Álvarez & Nicieza, 2002; Nicieza & Metcalfe, 1997) and, therefore, any observed costs of compensatory growth would not be caused by malnutrition, but by differential growth patterns (Lee et al., 2013). Hence, one vital cue with relevance for compensatory growth plasticity is temperature recognition. In the nematode worm *Caenorhabditis elegans*, the cold-sensitive TRP channel TRPA-1 (regulated by the *trpa-1* gene), can detect drop in temperature and regulate lifespan by modulating IIS signaling (Xiao et al., 2013). Furthermore, TRPA-1 acts in an age-and thermal-specific manner by extending lifespan upon adult cold-exposure but reducing it upon larval cold-exposure via signaling to DAF-16 in the IIS pathway, which in turn alters gene expression (Xiao et al., 2013; Zhang et al., 2015). The reduced lifespan under low larval temperature is very intriguing from a compensatory growth perspective, as it is a potential cost of compensatory growth after suboptimal juvenile conditions (Lee et al., 2013). In particular, Zhang et al., (2015) placed mature *C. elegans* adults to oviposit at 15°C, 20°C and 25°C and allowed the resulting offspring to develop in their respective temperature until the last larval instar, after which they were transferred to the optimal temperature of 20°C. They found that individuals reared at 15°C as juveniles lived shorter and did not pay any reproductive costs in terms of offspring number. Although growth was not measured in Zhang et al., (2015), their results are compatible with the model of costly compensatory growth after juvenile exposure to unfavorable conditions, regulated by *trpa-1* and the IIS pathway.

We set out to investigate the presence, costs and benefits of compensatory and slowed-down growth, as well as its genetic regulation, using the nematode *C. elegans*. In order to do so, we measured compensatory and slowed-down growth, age-specific reproduction and lifespan in wild-type worms as well as worms with a null-mutation in the *trpa-1* gene. We predict the *trpa-1* gene to be involved in regulation of compensatory growth and thus the null-mutation should result in absence of compensatory/slowed-down growth.

## Methods

### Strains

We used the wild-type strain N2 and the mutant *trpa-1(ok999)*, orthologue of human *TRPA-1*, which has a role in ion channel activity and is located in the IV chromosome (Kindt et al., 2007).

### Experimental design

Worms were reared on agar plates with NGM medium and antibiotics (streptomycin and kanamycin) and fungicide (nystatin) (Lionaki & Tavernarakis, 2013) and seeded with the antibiotic-resistant OP50-1 (pUC4K) *Escherichia coli* strain. Worms of both strains were revived from freezing, bleached twice and reared under standard laboratory conditions (20°C, 60% RH and no light) before the experiment to synchronize generation time and remove infections. We exposed newly laid eggs to one of three temperature treatments (Figure 1), under which the larvae would develop until reaching the third larval instar (L_3_), whereafter they were moved to a standardised temperature (20°C). The 20°C treatment is the normal temperature during standard laboratory conditions (and the most favourable temperature used in the study), and development to L_3_ takes 42h. The 15°C treatment delays development substantially, and it takes 72h to reach L_3_. This treatment is designed to induce compensatory growth, if present. Finally, the 25°C treatment speeds up development, and L_3_ is reached after 32h. The development time to L_3_ was the same for both strains and determined in pilot studies. The experiment was designed so that worms of all larval temperature treatments would simultaneously reach the L_3_, and therefore the treatment used different setup times. In order for maternal age effects not to influence the worms, all treatments were set up with different mothers for the different treatments, always of the same age (two days of adulthood, during peak reproduction). To setup the treatments, three plates were prepared for each strain and temperature combination, with 20 randomly selected mature adults ovipositing for 2h at 20°C in each plate. Once the synchronized ovipositions were terminated, adults were removed and the plates with the eggs were assigned to one of three temperature treatments: 15°C, 20°C or 25°C. Eggs were allowed to hatch and develop in the assigned temperature treatments until the achievement of the L_3_ stage (i.e. at 72, 42, 32 h of age for 15°C, 20°C and 25°C, respectively). When L_3_ was reached, 60 randomly selected worms per treatment per strain (N=360) were placed into individual 35 mm petri dishes and moved back to 20°C. Consequently, they were used to measure size, growth rate, lifetime reproductive success and fitness. From those 60 worms, the 40 worms with lowest ID labels in each strain and treatment (N=240) were photographed every 24h for five consecutive days for growth rate assessment starting from the day when L_3_ larvae were switched to 20°C (this constitutes day 0). Since the ID label was assigned at random, the selection of 40 out of the 60 worms in each strain and treatment combination constitutes a random selection in relation to any phenotype. Size was measured from photographs as the total area of the worm, using the software ImageJ (v1.46, Schneider et al. 2012). Growth was calculated as the size gained since the last measurement, divided by the time elapsed since last measurement. All worms were checked every 24h and, if still alive, moved to a new plate. The old plate was kept in order for eggs to hatch, and offspring were after 48h counted for daily fecundity assessment. The experiment was terminated when all focal worms had died.

**Figure 1.**
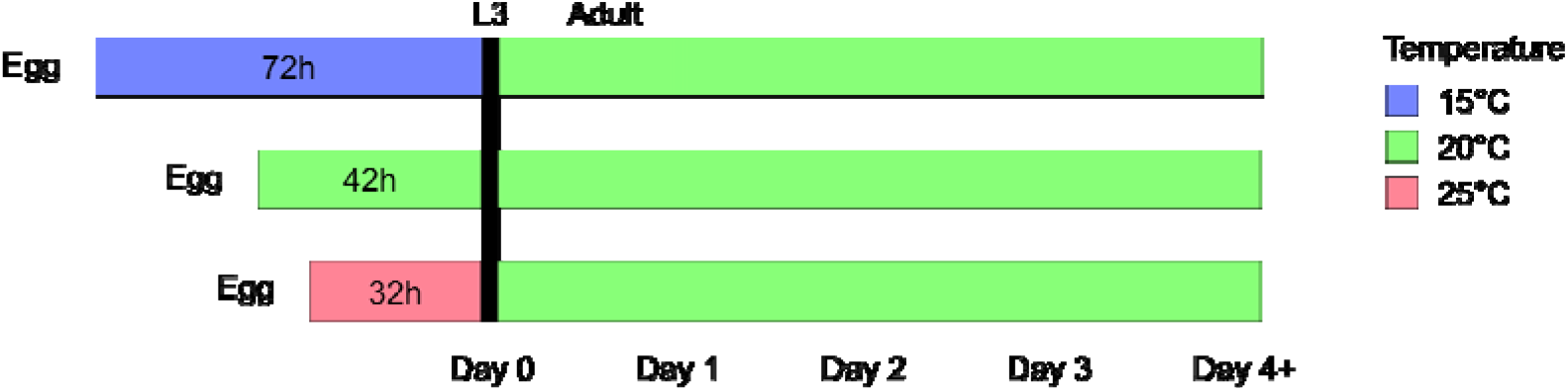
Experimental design. Overview of the experimental design. Eggs from parents reared in 20°C were placed into the three larval temperature treatments, where larvae developed until reaching the L_3_ larval stage. Thereafter, they were kept in 20°C for the remainder of the experiment. Juvenile treatments were synchronized so that L_3_ was reached at the same time in every treatment.

### Statistical analyses

All data were analyzed with *R 3*.*6*.*1* (R Core Team, 2014). Overall body size and growth rate were analyzed with a general linear mixed model approach for repeated measures implemented in the *lme4* package (Bates et al., 2015) including day (which is equivalent to an age counted in days), larval temperature treatment, strain and all two-and three-way interactions as fixed effects. We included worm ID as a random intercept and a random slope effect to account for individual variation in area or growth rate with age. Furthermore, we also included day^2^ as fixed effect and all interactions with day^2^ to account for curvature. We deleted in a step-wise process non-significant, 3-way interaction terms and present the model with the smallest AIC value. Significance of fixed parameters were assessed using χ^2^ tests implemented in the *car* package (Fox & Weisberg, 2010). For analyses of growth, we also included the initial size at the start of each growth interval as a covariate, to control for any size-dependency of growth.

Reproduction was analyzed both as lifetime reproductive success (defined by the number of hatched offspring) and as two rate-sensitive measure of reproduction, which also takes the timing of reproduction into account (see below). These rate-sensitive measures are analogous to the intrinsic rate of population growth (Brommer et al., 2002). As rate-sensitive measures, they give increased weight to early reproduction and short development time. We calculated these rate-sensitive measures by constructing a population projection matrix for each individual, and then calculated the dominant eigenvalue from this matrix (McGraw & Caswell, 1996).

We calculated the rate-sensitive measures (1) *individual fitness* λ_*ind*_ and (2) *development-independent reproductive rate*, which only differs in how development-time was incorporated. For *individual fitness* λ_*ind*_ we included the treatment-exposure times in the calculation of development time (rounded to whole days, as this was the precision of the reproductive data). In our experimental design, we synchronized the emergence of our focal individuals as L_3_ larvae, by rearing them to L_3_ at three different temperatures (Figure 1). Therefore, analysing individual fitness investigates whether the worms can fully compensate for the substantial fitness loss caused by the long development time in cold juvenile temperature. In addition, we also calculated *development-independent reproductive rate* as the rate of egg production after L_3_, without including the temperature-induced differences in the time required to attain L_3_. This metric uses a standardized development time for all treatments in the projection matrix (2 days). The reason for calculating this metric is to investigate if the reproductive schedule has been altered.

Lifetime reproductive success, individual fitness λ_ind_ and reproductive rate were investigated with a general linear model approach including larval temperature treatment, strain (*N2* or *trpa-1*) and their interaction as fixed effects. Finally, we tested lifespan with a Cox proportional hazard regression model (package *survival*) including larval treatment, strain and their interaction as fixed effects. Lifespan was measured as *lifespan since treatment end* by setting L_3_ stage as day 0, in order to separate any effects of the treatments from the direct effects of the developmental delay caused by the treatments. This corresponds to adult lifespan since the first day of adulthood is day 1. However, as for rate-sensitive reproductive measures, we also analysed the data including the different development times directly caused by the treatment (total lifespan from egg). Worms dying of matricide (internal hatching of eggs) were censored.

Five individuals remained small throughout the whole experiment, all in the 15°C/*trpa-1* treatment (these individuals are highlighted in figure S1). As our focus was on growth rate, these individuals were excluded from all main analyses, although their exclusion did not alter the result (supplementary tables S8-S11).

Since the *trpa-1* worms in 20°C showed an aberrant growth pattern, all analyses were performed both with the full dataset and with a reduced dataset, where the 20°C treatment was removed for both strains. This was to ensure that the *trpa-1* in 20°C was not driving any pattern found.

## Results

We found that both strain and larval temperature treatment affected size, and these factors interacted with day and day^2^ (Table 1, Figure 2). AIC model selection indicated that the best model for body size was the one without any of the three-way interactions. Body size increased significantly with day, but treatment effects differed between the strains. For growth rate, AIC model selection showed that the best model also included the three-way interactions with day and day^2^ (Table 2, Figure 3). The 15°C treatment group in the wild-type strain responded to the switch to 20°C by accelerating growth and growing faster than the 20°C treatment (it showed compensatory growth). As a result, they fully caught up in size and reached the same final size as the worms from the 20°C treatment. In contrast, the 25°C treatment had the opposite response; growth decelerated and was slower than the 20°C treatment, which resulted in worms never reaching the size of individuals from the 20°C treatment. Overall, growth rate decreased with age showing a strong interaction between strain and treatment (Table 2).

**Table 1.** Size. The effect of juvenile temperature treatment (15°C, 20°C and 25°C), strain (wild-type, *trpa-1*), day, day^2^ and their interactions on the daily size (measured as area). This final model was obtained by model simplification using AIC.

| Parameter | $\chi^2$ | d.f. | p |
| --- | --- | --- | --- |
| (Intercept) | 477.16 | 1 | <0.001 |
| Treatment | 57.32 | 2 | <0.001 |
| Strain | 4.25 | 1 | 0.039 |
| Day | 3500.34 | 1 | <0.001 |
| Day<sup>2</sup> | 2110.56 | 1 | <0.001 |
| Treatment × Strain | 40.01 | 2 | <0.001 |
| Treatment × Day | 7.58 | 2 | 0.023 |
| Strain × Day | 7.40 | 1 | 0.007 |
| Treatment × Day<sup>2</sup> | 13.07 | 2 | 0.001 |
| Strain × Day<sup>2</sup> | 18.46 | 1 | <0.001 |

**Table 2.**
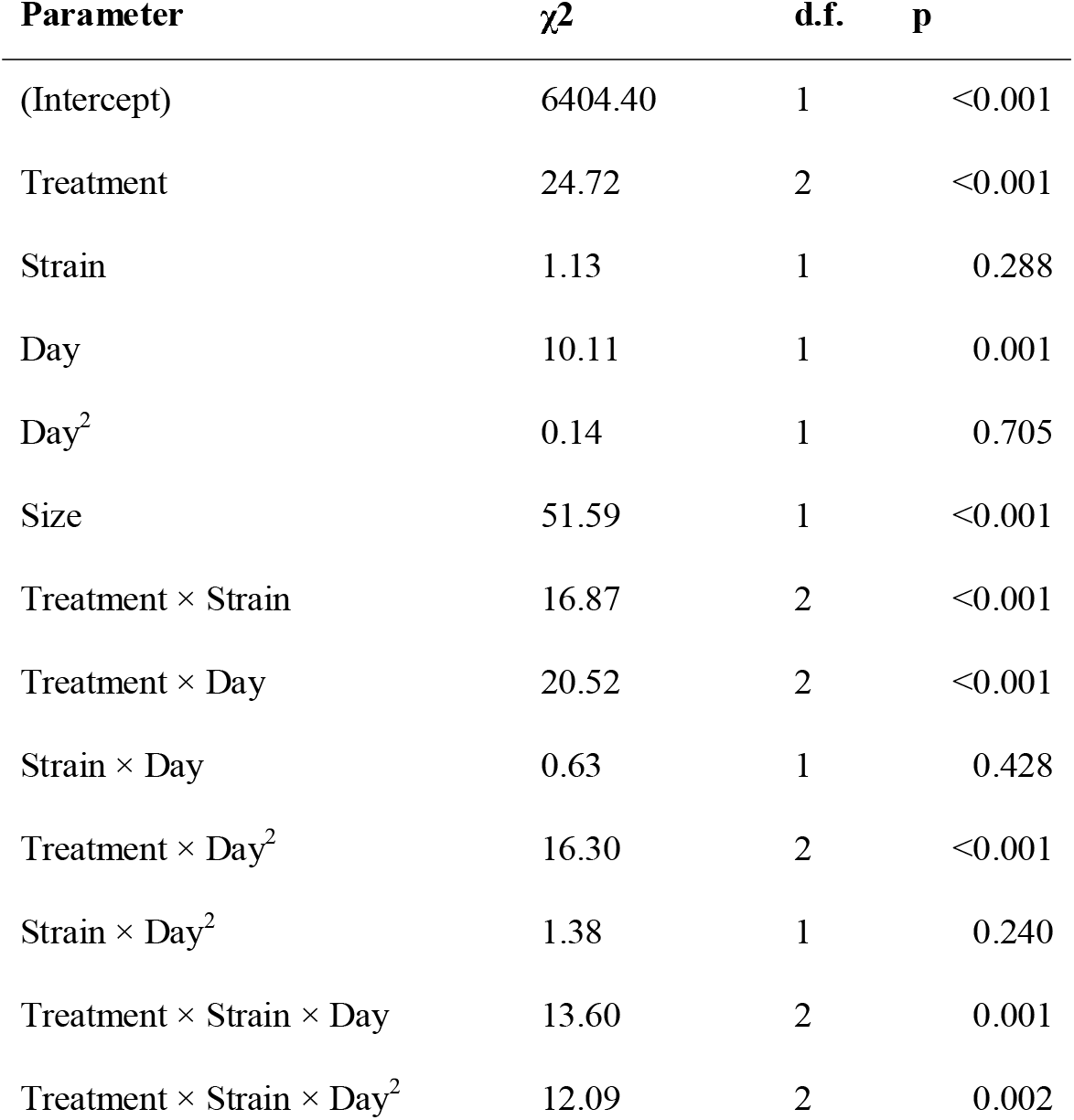
Growth rate. The effect of juvenile temperature treatment (15°C, 20°C and 25°C), strain (wild-type, *trpa-1*), day, day^2^ and their interactions on the daily growth rate, with Size at the start of each day included as a covariate to control for size-specific growth rate.

**Figure 2.**
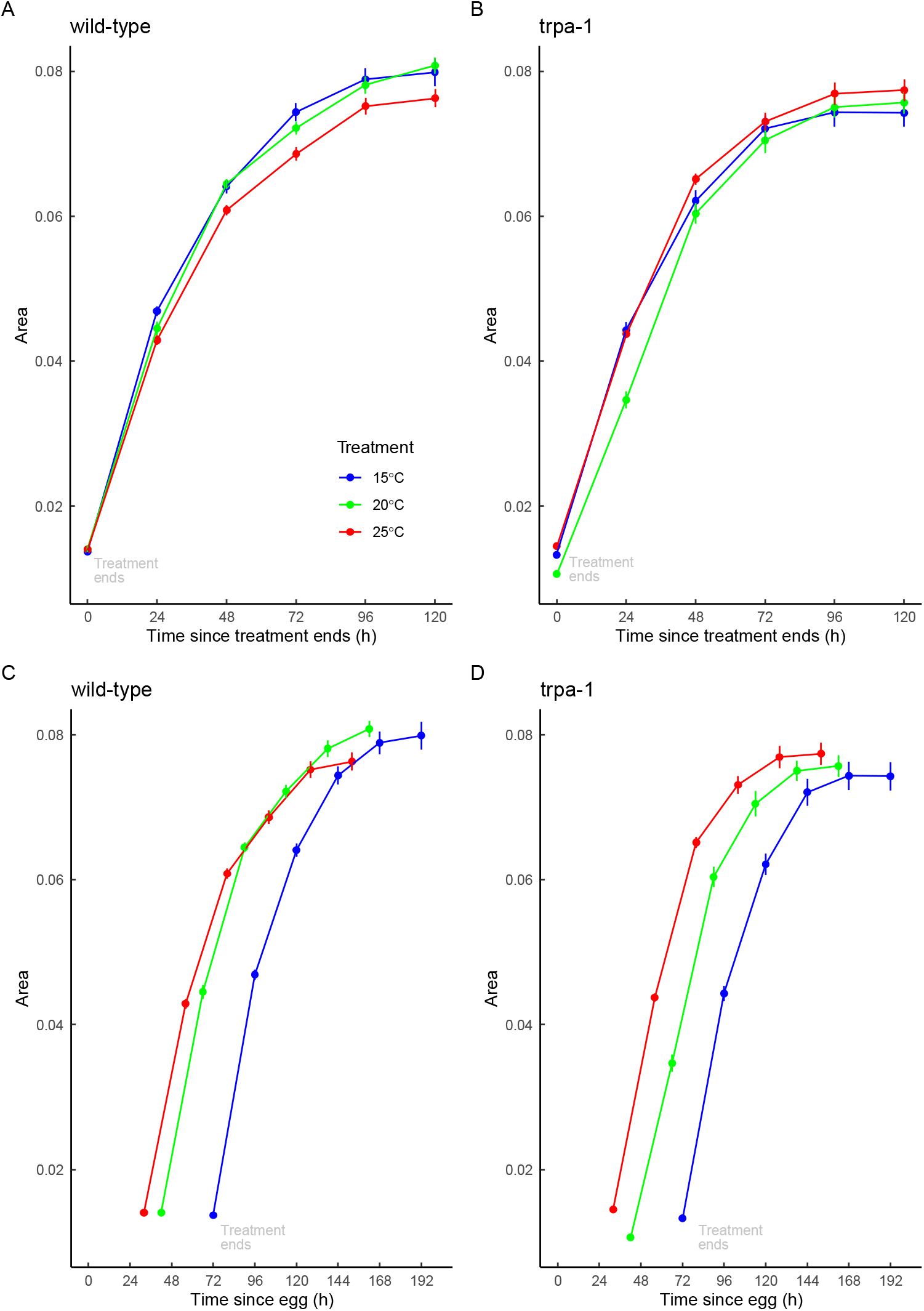
Variation in body size measured as body area in mm^2^ from the end of the larval treatment for (A) wild-type and (B) *trpa-1* mutant worms. In order to illustrate the developmental delay caused by the juvenile treatment, the same data is also presented using time since egg for (C) wild-type and (D) *trpa-1 mutants*. End of the juvenile treatment (day 0) is the day on which the switch to 20°C was performed, corresponding to the molt from L_2_ to L_3_. In blue, green and red are displayed 15°C, 20°C and 25°C larval treatments respectively. Symbols represent mean ± SE.

**Figure 3.**
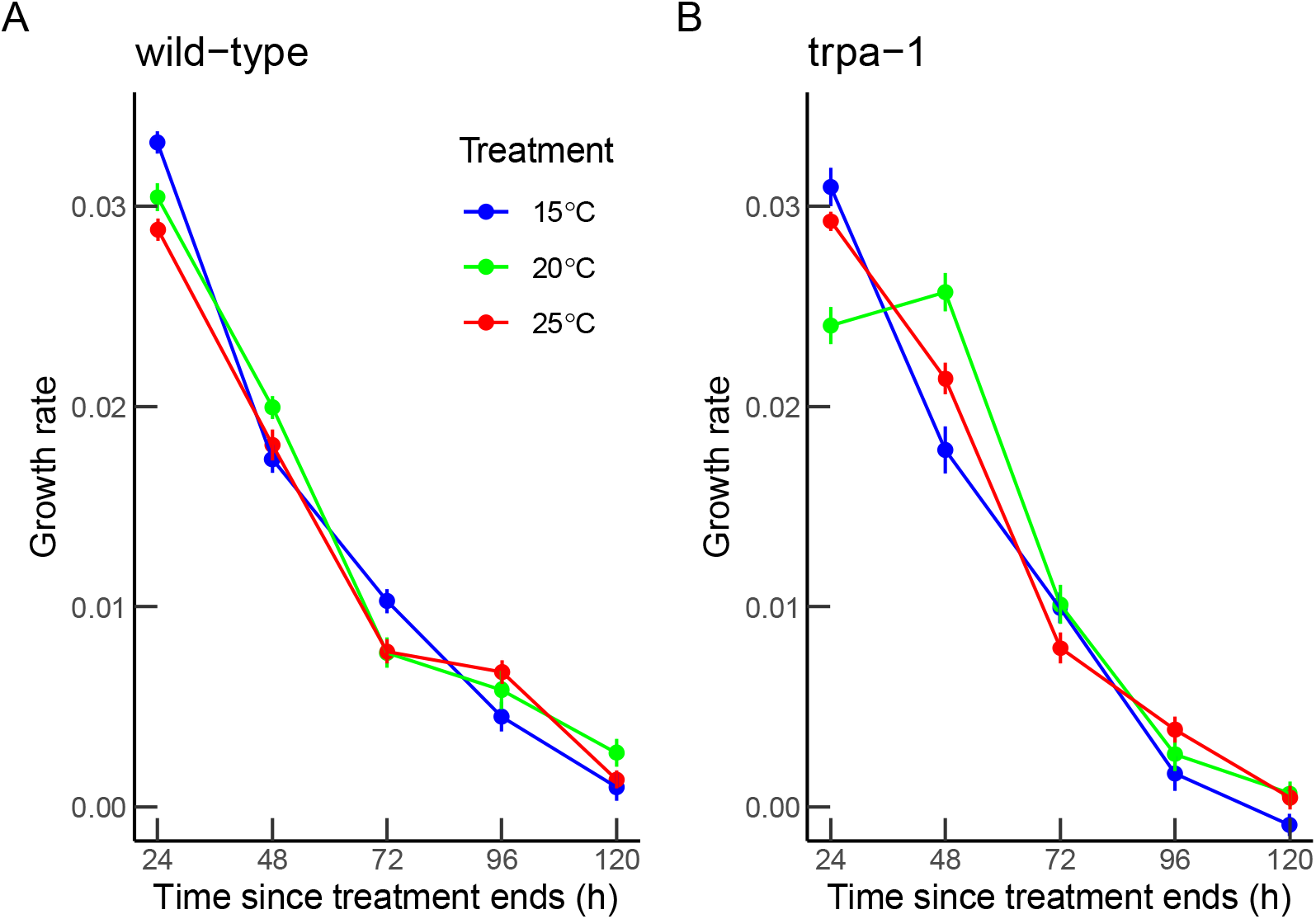
Daily growth rate since the end of the juvenile treatment for (A) wild-type and (B) *trpa-1* mutant worms. In blue, green and red are displayed 15°C, 20°C and 25°C larval treatments, respectively. Symbols represent mean ± SE.

There was little evidence of compensatory growth in the *trpa-1* mutant worms (Figure 3). However, the *trpa-1* mutant worms from the 20°C treatment showed a very small size at L_3_ and thus differed strongly from the 20°C wild-type treatment worms. To ensure that the interaction between strain and treatment was not driven by the *trpa-1* and 20°C combination, we also performed this and all other analyses of all traits both with and without the 20°C treatment (for both strains). When removing the 20°C treatment from the models, we still found a strong interaction between strain and treatment, reinforcing our interpretation of compensatory growth only being present in the wild-type (Supplementary table 2).

We used the age-specific reproduction data (Figure 5) to analyze individual fitness λ_ind,_ development-independent reproductive rate and lifetime reproductive success. When analysing individual fitness, which was calculated using the treatment-specific development times, we found highest fitness in the juvenile treatment that was applied for the shortest duration (25°C) and lowest fitness in the most long-lasting juvenile treatment (15°C) (Figure 4B, supplementary table S6). We also found an interaction between strain and treatment, caused by comparably lower fitness for *trpa-1* in 15°C.

**Figure 4.**
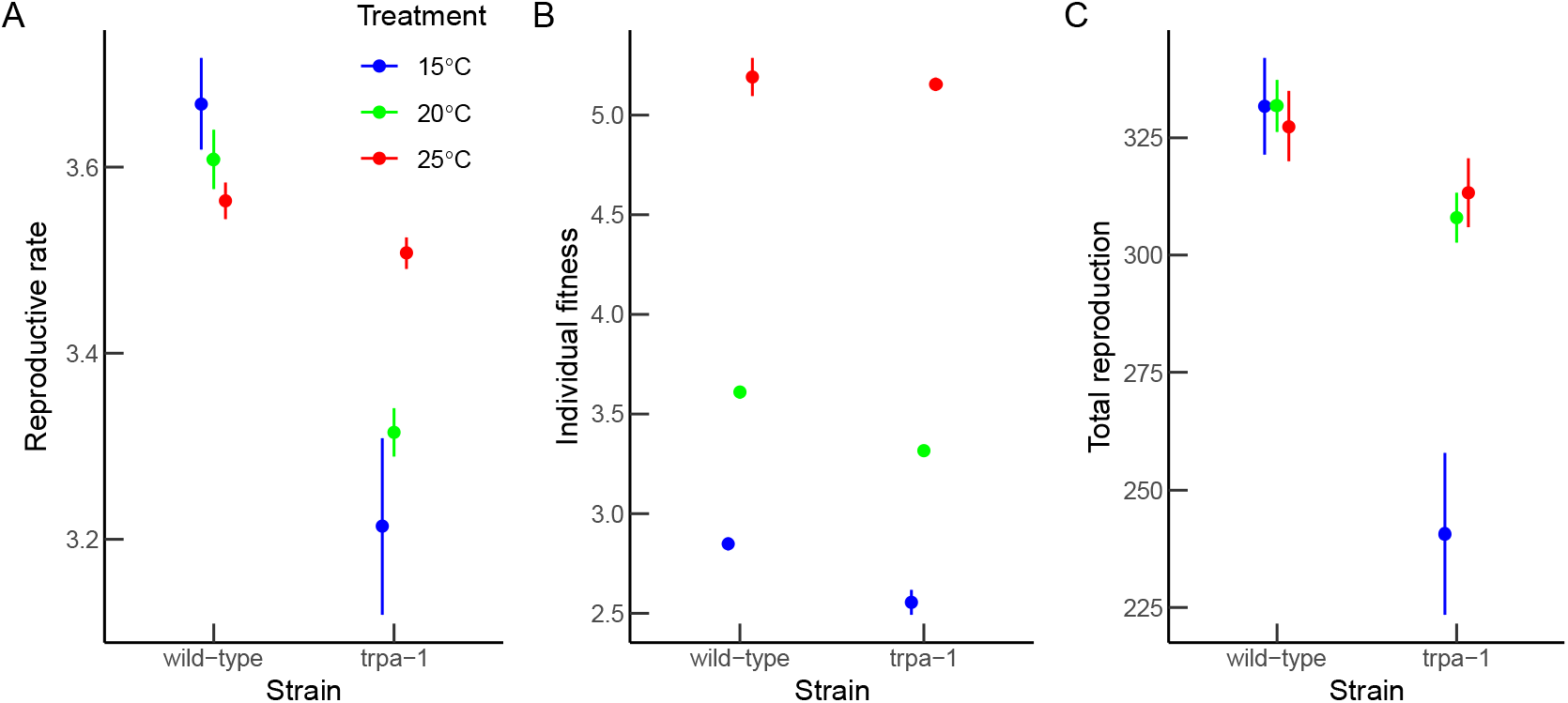
(A) Development-independent reproductive rate, (B) individual fitness λ_ind_ and (C) lifetime reproductive success for wild-type and *trpa-1* mutant worms. In blue, green and red are displayed 15°C, 20°C and 25°C juvenile treatments, respectively. Symbols represent mean ± SE.

**Figure 5.**
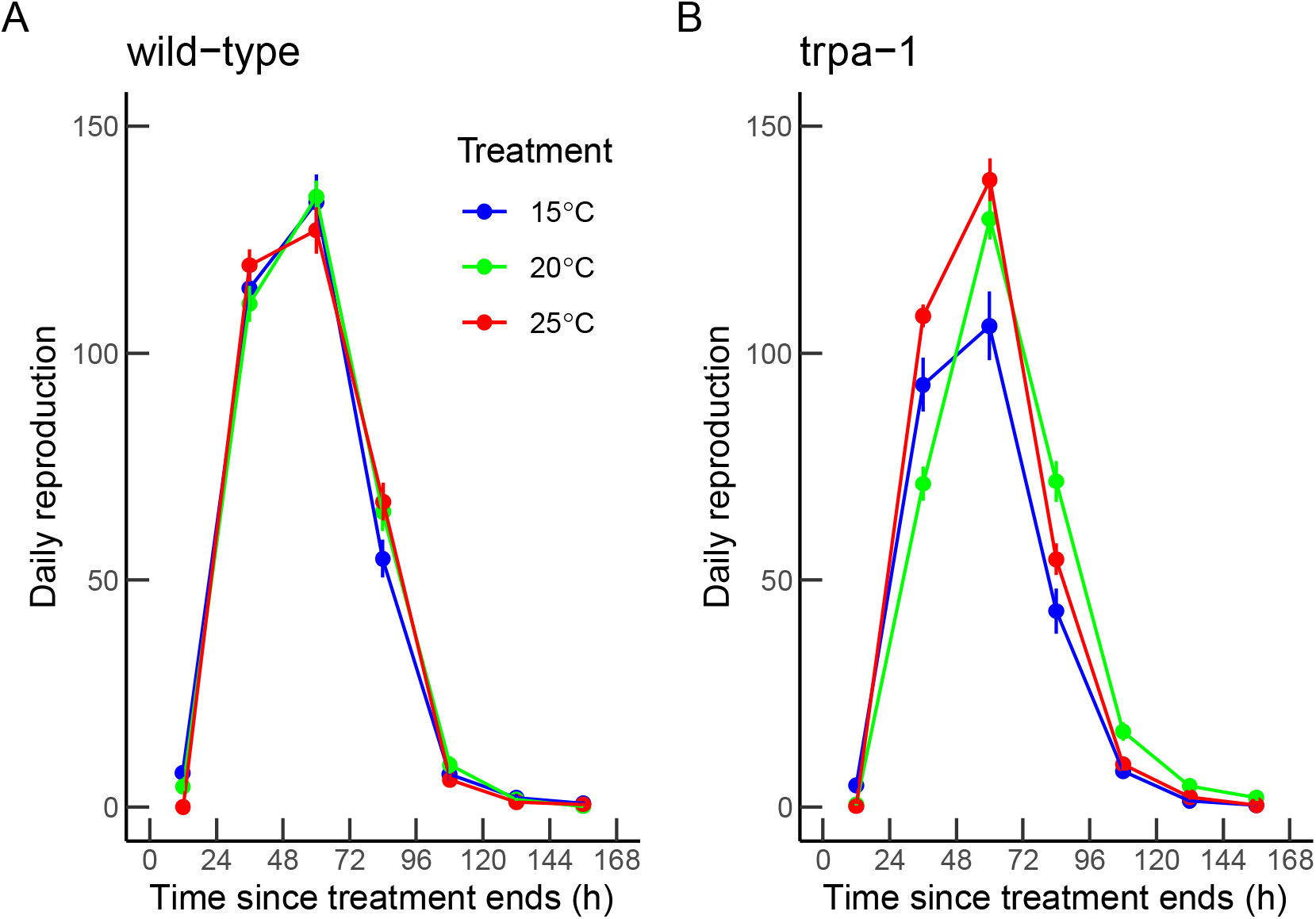
Daily reproduction for (A) wild type and (B) *trpa-1* mutant worms. In blue, green and red are displayed 15°C, 20°C and 25°C juvenile treatments, respectively. Symbols represent mean ± SE.

For development-independent reproductive rate, which investigates changes in reproductive rate after the end of the juvenile treatment, we found a significant interaction between strain and treatment (Table 3, Figure 4A). While the reproductive rate was generally higher in wild-type worms, the ordering of the treatment differed in the two strains; the 15°C wild-type worms tended to have a higher rate of offspring production after maturity than the wild-type 20°C worms. The 15°C *trpa-1* mutants did now show such compensation and in fact have a lower rate of offspring production and lower lifetime reproductive success. Thus, although all worms exposed to 15°C have low individual fitness because of a very long development time, the wild-type worms partly compensated by increasing their reproductive rate after being released from the low juvenile temperature. In contrast, in *trpa-1* mutants the 15°C treatment had much lower fitness, as well as larger variance in fitness, caused by several worms hardly reproducing. Lifetime reproductive success was also higher in wild-type worms but did not differ among the three temperature treatments. The 15°C treatment thus shifted when they laid eggs, but did not change the total number of offspring produced (Table 3, Figure 4C). Instead, the interaction was only driven by the *trpa-1* mutants, where lifetime reproductive success was lowest for worms that developed at 15°C. Again, removing the 20°C treatment did not alter the main findings for individual fitness or lifetime reproductive success (Supplementary table 3).

**Table 3.** Reproductive rate and lifetime reproductive success. The effect of juvenile temperature treatment (15°C, 20°C and 25°C), strain (wild-type, *trpa-1*), and their interactions on reproductive rate and lifetime reproductive success.

| Reproductive rate |  |  |  |  |  | Lifetime reproductive success |  |  |  |  |
| --- | --- | --- | --- | --- | --- | --- | --- | --- | --- | --- |
| Parameter | d.f. | SS | MS | F | p | d.f. | SS | MS | F | p |
| Treatment | 2 | 0.32 | 0.16 | 1.35 | 0.262 | 2 | 47259 | 23630 | 4.52 | 0.012 |
| Strain | 1 | 4.76 | 4.76 | 40.59 | <0.001 | 1 | 119372 | 119372 | 22.84 | <0.001 |
| Treatment × Strain | 2 | 1.48 | 0.74 | 6.32 | 0.002 | 2 | 61207 | 30603 | 5.86 | 0.003 |
| Residuals | 343 | 40.22 | 0.12 |  |  | 343 | 1792742 | 5227 |  |  |

Despite the presence of compensatory growth in wild-type worms, we did not find any lifespan cost when lifespan was calculated from the end of the juvenile treatment, since adult lifespan did not differ between strains or treatments (Figure 6, supplementary tables S4-S5). However, when including the different treatment durations in the analysis of total lifespan from egg, we found the longest lifespan in the 15°C treatment, and shortest lifespan in the 25°C treatment (supplementary figure S2, supplementary table S7), which reflects the longer development time in 15°C and shorter development time in 25°C.

**Figure 6.**
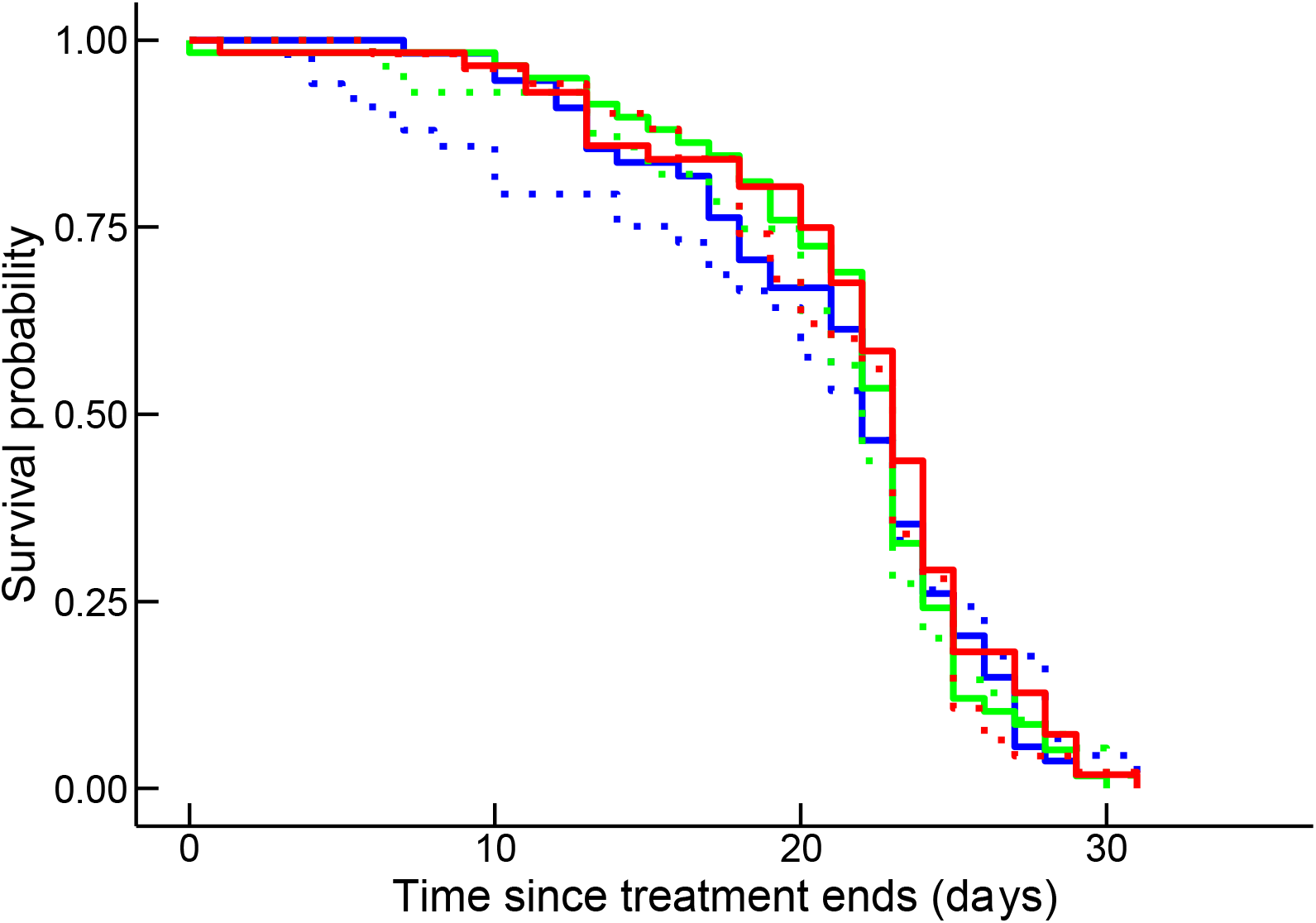
Lifespan, starting from the end of the larval treatment, which is different from total lifespan since the differences among treatment groups in the age at L_3_ is not included. Solid and dotted lines indicate wild-type and *trpa-1* mutants, while blue, green and red indicate 15°C, 20°C and 25°C juvenile treatments.

## Discussion

We found that wild-type *C. elegans* nematodes express compensatory growth as a response to unfavorable juvenile conditions. In particular, we found that larvae of wild-type worms exposed to low temperature of 15°C slowed down their growth and were delayed in time as they needed more than 70% longer to reach a similar body size as worms from standard conditions (72 h vs. 42 h respectively) (Figure 1). This delay in growth translated into equivalent delay in transition into L_3_, since *C. elegans* needs to reach a certain threshold size to moult to the next larval stage (Uppaluri & Brangwynne, 2015). However, once conditions returned to a more favourable temperature, they compensated for the larval cold exposure and reached the final adult size equivalent to the size of unrestricted individuals (catch-up growth). This follows a common observation that organisms, after being released from suboptimal juvenile conditions, show compensatory growth to compensate for the bad start (Hector & Nakagawa, 2012; Jobling, 2010).

Lifetime exposure of *C. elegans* to 15°C has been shown to result in decreased lifetime reproductive success (Gouvêa et al., 2015) suggesting that 15°C is a suboptimal environment. Organisms born in suboptimal conditions are expected to perform worse and have lower fitness than those born in optimal conditions (Monaghan, 2008) and this is especially true if the developmental environment is mismatched with the adult environment (Monaghan, 2008) as in our case. Interestingly, once worms returned to favourable conditions (20°C) they were able to partly mitigate the negative carry-over effect of suboptimal juvenile temperature and reached lifetime reproductive success comparable to reproduction under standard conditions. In addition, although the extended development in the cold juvenile treatment reduced the individual fitness, once released from the suboptimal juvenile treatment they increased their development-independent reproductive rate with no apparent cost in adult lifespan. The increased reproductive rate was likely caused by higher reproduction during the first day of adulthood, since rate-sensitive reproductive measures are very sensitive to early reproduction (Brommer et al., 2002).

Larval exposure to high temperature (25°C) resulted in faster juvenile growth as worms reached the same larval size and L_3_ transition faster compared to those raised in 20°C. However, after the switch to 20°C, worms decreased their growth, which resulted in smaller final body size. Decreased growth mirrors a previous finding in sticklebacks (Lee et al., 2013) and may reflect a cost of accelerated larval growth or a cost of exposure to higher temperatures during larval development. Their lifetime reproductive success and reproductive rate was also comparable to the 20°C control.

Since compensatory growth is expressed as a response to the developmental delay caused by bad growth conditions early in life, organisms are expected to show a stronger response the more they are time constrained (Orizaola et al., 2014). If they manage to increase growth after returning to better conditions, this may result in trade-offs with lifespan or reproduction. These types of trade-offs are notoriously difficult to measure experimentally, since it would ideally be done by designing an experiment where a random sub-set of individuals perform compensatory growth, while other individuals do not. Our approach to compare treatments where compensatory growth is present or absent are thus necessarily confounding treatment effects with effects of compensatory growth. Nevertheless, this approach can still provide insights in whether any trade-offs are correlated with the presence of compensatory growth.

Surprisingly, we did not find that adult lifespan or total reproduction was lower in the 15°C juvenile treatment than in the 20°C, despite the expression of compensatory growth. In fact, worms from the 15°C juvenile treatment managed to increase their development-independent reproductive rate. It should however be noted that this increase in reproductive rate could only partially compensate for the substantial fitness loss caused by the increased development time in 15°C. Compensatory growth could therefore not restore individual fitness to that of the 20°C treatment.

Since individual fitness is very sensitive to differences in development time (Brommer et al., 2002), the direct effect of the duration of the juvenile treatments explained all differences in individual fitness between treatments, with highest fitness values for the 25°C treatment, and lowest for the 15°C treatment. This does not imply that the warm environment is the optimal environment, only that 25°C shortens the development time the most. In fact, lifelong exposure to 25°C increases individual fitness but reduces total reproduction compared to 20°C in the sister species *C. remanei*, because of the development time effect (Lind et al., 2020). Similarly, the total lifespan is also explained by the direct effects of the juvenile treatment, where the long juvenile exposure to 15°C gives this treatment a longer total lifespan. This does not mean that compensatory growth extends lifespan. Instead, lifespan measured from the end of the juvenile treatment (from L_3_), which is equivalent to adult lifespan (since all worms matured during day 1), is the same for all treatments.

While compensatory growth is found across several taxa, it is not always present, and can instead be seen as a strategy that is adaptive mainly in time-constrained and seasonal environments, or when large size is under strong selection. For example, compensatory growth is present in amphibians from time-constrained northern latitudes, but not in southern populations (Dahl et al., 2012; Orizaola et al., 2014). Similarly, sticklebacks from ponds that are selected for large size show compensatory growth, but the smaller marine sticklebacks do not (Ab Ghani & Merilä, 2015).

Since *Caenorhabditis* nematodes have reduced female reproduction if adult body size is pharmacologically reduced (Lind et al., 2016), and the natural habitat of *C. elegans* is ephemeral with rapidly decaying plant material (Félix & Braendle, 2010) creating strong time-constraints, compensatory growth is, therefore, predicted to be adaptive under natural settings. Therefore, it is not surprising that individuals that were as juveniles raised in 15°C, which are delayed in time as they switched to L_3_ later then worms raised at warmer temperatures, will be under pressure to ‘make up for the lost time’. The observation that, although delayed in time, worms subjected to 15°C were not smaller at the shift to L_3_, is probably explained by the fact that *C. elegans* needs to reach a threshold size to transition between larval stages (Uppaluri & Brangwynne, 2015).

Although compensatory growth is common across taxa (Ab Ghani & Merilä, 2015; Burraco et al., 2020; Fisher et al., 2006; Lee et al., 2013; Morgan & Metcalfe, 2001; Murillo□Rincón et al., 2017; Orizaola et al., 2014; Ozanne & Hales, 2004; Royle et al., 2006), its molecular regulation remains unknown. Here, we show that compensatory growth in response to low juvenile temperature is regulated in part by the thermo-sensitive TRPA-1 ion channel in *C. elegans*. While wild-type worms respond to low juvenile temperature with compensatory growth once returned to favorable temperature, the *trpa-1* knockout mutants did not increase their growth rate. Furthermore, they also did not show any associated response to temperature treatment in reproductive rate, in contrast to the wild-type individuals.

This constitutes, to our knowledge, the first example of a gene involved in the regulation of compensatory growth plasticity. TRPA-1 ion channel acts as an upstream regulator of DAF-16 in the insulin-like signaling (IIS) pathway (Xiao et al., 2013; Zhang et al., 2015). As the only FOXO homolog in *C. elegans*, DAF-16 integrates upstream signaling and directs multiple responses in life-history and physiology, such as development, stress resistance and ageing, and DAF-16/FOXO is conserved from worms to mammals (reviewed in Zečić & Braeckman, 2020). Moreover, since the IIS pathway integrates multiple environmental cues (Feeney et al., 2016; Metaxakis et al., 2014; Sim et al., 2015) and induces plastic responses across organisms (Green & Extavour, 2014; Mortzfeld et al., 2019; Snell-Rood & Moczek, 2012) it has been suggested as a master regulator for adaptive phenotypic plasticity (Regan et al., 2020). Our finding that TRPA-1, which integrates into the IIS pathway, is involved in the regulation of compensatory growth plasticity in response to temperature in *C. elegans* reinforces this view. Since the IIS pathway regulates multiple life-history traits in *C. elegans* (Zečić & Braeckman, 2020) it is not surprising that the effects of the *trpa-1* knockout mutants are not limited to compensatory growth, but also result in an overall lower egg production across treatments and influence the ranking of growth rate and fitness among treatment groups.

Surprisingly, while we found reduced fitness by exposure to low temperatures, we found no other life-history cost correlated with compensatory growth, neither in total reproduction nor in adult lifespan. For reproduction, we instead found that worms experiencing compensatory growth maintained the same total reproduction and in addition had an increased reproductive rate when standardizing development times, caused by an earlier reproductive peak. Our finding that the lifetime reproductive success was unaffected by the knockout of *trap-1* mirrors the previous findings of Zhang et al. (Zhang et al., 2015), but we additionally show that further insight into the reproductive performance is gained by using rate-sensitive measures of reproduction, that takes timing of reproduction into account (Brommer et al., 2002; McGraw & Caswell, 1996). Since the natural habitat of *C. elegans* is decaying plant matter, populations show a rapid boom and burst reproductive strategy (Frézal & Félix, 2015), and thus, the importance of early reproduction is eminent.

While reproductive consequences of compensatory growth are seldom measured, the few studies with such data have mixed findings. For instance, Dmitriew & Rowe, (2007) detected no reproductive consequences in ladybird beetles, while Auer et al., (2010) found that female guppies experiencing compensatory growth had lower offspring production.

Our findings that rate-sensitive measures of reproduction are important suggest that future studies should focus not only on viable offspring number but also integrate the timing of reproduction into rate-sensitive measures such as reproductive rate and individual fitness (Brommer et al., 2002). It should be, however, noted that similarly to previous studies we did not measure offspring quality, but only the number of offspring that survived for at least 48 hours (i.e. until the reproduction measurement was taken). The potential trade-off between number and quality of offspring is well known in the life history literature (Roff, 1993; Stearns, 1992), and such inter-generational trade-offs have recently been found in *Caenorhabditis* nematodes (Mautz et al., 2020) but are not ubiquitous (Lind et al., 2019).

While reproductive consequences of compensatory growth are under-studied, its consequences for lifespan have been in greater focus. A number of studies now provide evidence for lifespan costs across taxa (Inness & Metcalfe, 2008; Lee et al., 2013), even if costs in lifespan are sometimes only present under certain environmental conditions (Dmitriew & Rowe, 2007). In this study, we did not detect any costs in terms of adult lifespan (measured from L_3_ onwards), and we were, thus, unable to replicate the findings of Zhang et al., (2015), who found that cold-exposure during the juvenile period shortens lifespan in wild-type but not in *trpa-1* mutants. A possible explanation for the difference between the studies might be the use of the antibiotic kanamycin, which was used to fight bacterial infections and which was not used by Zhang et al., (2015). Kanamycin may potentially interact with the temperature treatment and affect the lifespan of worms. The two studies have relatively similar sample sizes per treatment, with 30 reproducing worms in Zhang et al., (2015) and 60 in the present work. Other possible costs of compensatory growth may not involve life-history traits, but can be detrimental for the organism under challenging natural conditions, such as impaired cognitive performance (Fisher et al., 2006), reduced immune-response (Murillo□Rincón et al., 2017) or higher antioxidant activities (Burraco et al., 2020).

## Conclusions

To conclude, we find that *C. elegans* express compensatory growth in response to suboptimal juvenile temperature, and that the cold-sensitive TRP ion channel TRPA-1, which is an upstream regulator of the IIS signaling pathway, regulates this response. These findings open for future studies of an ecologically relevant and widespread form of phenotypic plasticity using the powerful *C. elegans* model system and the associated phenotypic and molecular tools.

## Author contributions

ZS, ER, FS and ML designed the experiment, ZS, ER, FS, PIZ and ML performed the experiment, ER and PIZ measured the worms, ER and ML analyzed data, ZS, ER and ML drafted the manuscript. All authors contributed to revision of the manuscript and gave final approval for publication.

## Acknowledgements

We thank Andrea Hinas for aid with experimental design. This work was supported by the Swedish Research Council VR Grants 2016–05195 and 2020-04388 and the Carl Tryggers Stiftelse grant CTS 17: 285 to M.I.L. Some strains were provided by the CGC, which is funded by NIH Office of Research Infrastructure Programs (P40 OD010440).

## Data Accessibility Statement

Data is available from the Figshare respiratory: https://doi.org/10.6084/m9.figshare.14459964.v3 (Sekajova et.al, 2021).

