## Supplementary material for "Temperature-induced compensatory growth in *C. elegans* is regulated by a thermosensitive TRP channel and influences reproductive rate"

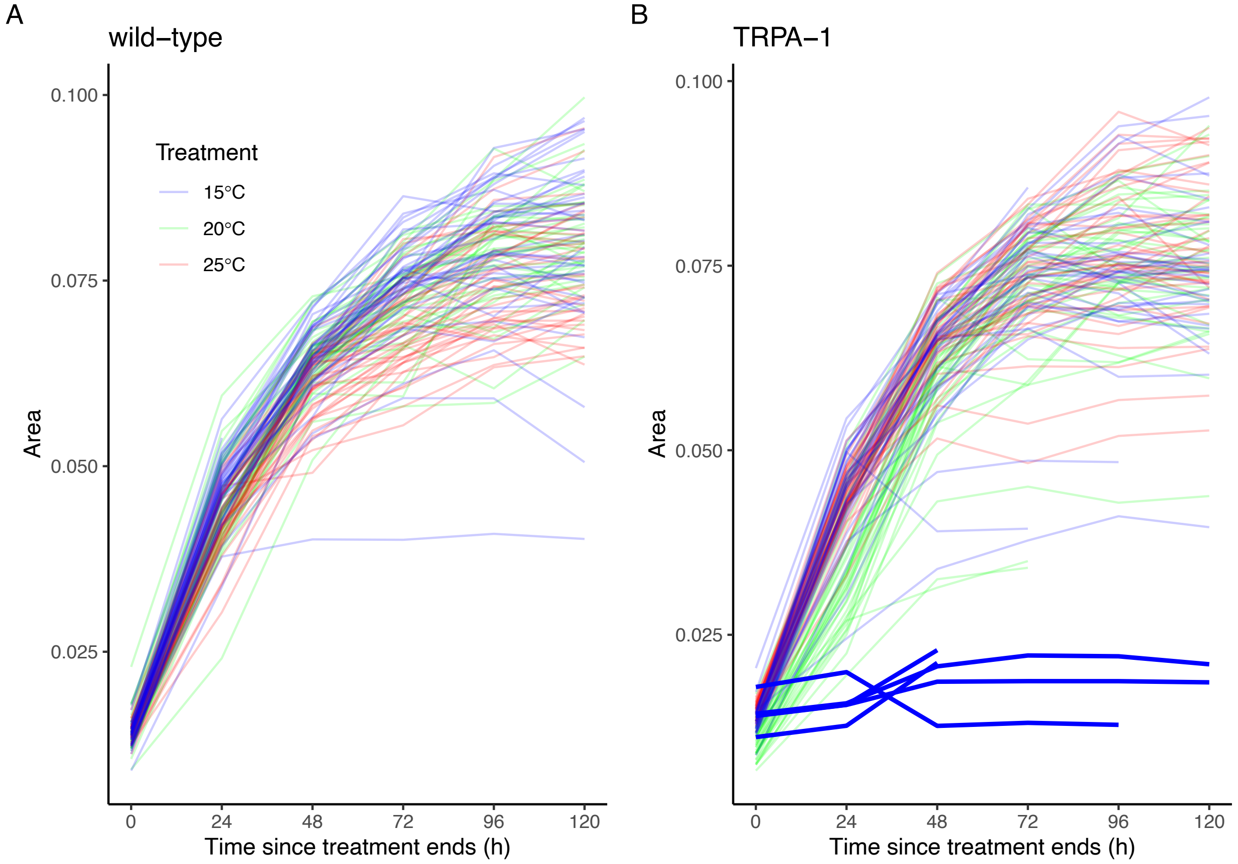


**Figure S1.** Variation in body size measured as body area in $mm^2$ from the end of the juvenile treatment for individual (A) wild-type and (B) *trpa-1* mutant worms. The five outlier worms that were excluded from all analyses because of a lack of growth are highlighted in bold.

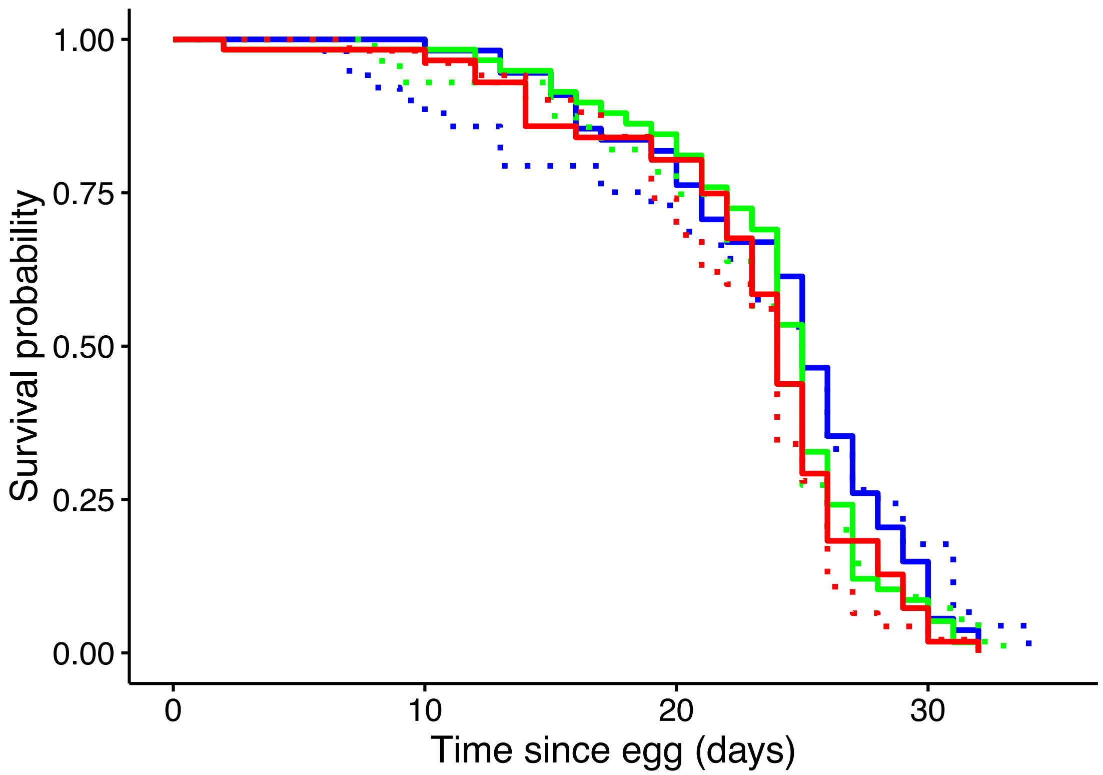


**Figure S2.** Lifespan with the differential length of the juvenile treatment taken into account. Solid and dotted lines indicate wild-type and *trpa-1* mutants, while blue, green and red indicate 15°C, 20°C and 25°C larval treatments.

**Table S1. Size excluding 20°C.** The effect of juvenile temperature treatment (15°C, 25°C), strain (wild-type, *trpa-1*), Day, Day$^2$ and their interactions on the daily size (measured as area). This final model was obtained by model simplification using AIC.

| **Parameter** | **χ2** | **d.f.** | **p** |
|---|---|---|---|
| (Intercept) | 31519.38 | 1 | <0.001 |
| Treatment | 8.95 | 1 | 0.003 |
| Strain | 3.85 | 1 | 0.050 |
| Day | 1971.97 | 1 | <0.001 |
| Day$^2$ | 1035.28 | 1 | <0.001 |
| Treatment × Strain | 9.70 | 1 | 0.002 |
| Treatment × Day | 4.71 | 1 | 0.030 |
| Strain × Day | 0.54 | 1 | 0.464 |
| Treatment × Day$^2$ | 4.40 | 1 | 0.036 |
| Strain × Day$^2$ | 1.01 | 1 | 0.314 |

**Table S2. Growth rate excluding 20°C.** The effect of juvenile temperature treatment (excluding 20°C), strain (wild-type, *trpa-1*), Day, Day$^2$ and their interactions on the daily growth rate, with Size included as a covariate to control for size-specific growth rate.

| **Parameter** | **$\chi$2** | **d.f.** | **p** |
|---|---|---|---|
| (Intercept) | 14375 | 1 | <0.001 |
| Treatment | 1.08 | 1 | 0.299 |
| Strain | 0.02 | 1 | 0.883 |
| Day | 17.58 | 1 | <0.001 |
| Day$^2$ | 1.65 | 1 | 0.198 |
| Size | 35.91 | 1 | <0.001 |
| Treatment × Strain | 6.51 | 1 | 0.011 |
| Treatment × Day | 8.89 | 1 | 0.003 |
| Strain × Day | 9.56 | 1 | 0.002 |

**Table S3. Individual fitness and total reproduction excluding 20°C**. The effect of juvenile temperature treatment (15°C, 25°C), strain (wild-type, *trpa-1*), and their interactions on individual fitness and total reproduction.

| | Individual fitness | | | | | Total reproduction | | | | |
|---|---|---|---|---|---|---|---|---|---|---|
| **Parameter** | **d.f.** | **SS** | **MS** | **F** | **p** | **d.f.** | **SS** | **MS** | **F** | **p** |
| Treatment | 1 | 0.10 | 0.10 | 0.67 | 0.413 | 1 | 36266 | 36266 | 5.19 | 0.024 |
| Strain | 1 | 2.40 | 2.40 | 15.84 | <0.001 | 1 | 110602 | 110602 | 15.84 | <0.001 |
| Treatment × Strain | 1 | 1.32 | 1.32 | 8.75 | 0.003 | 1 | 53485 | 53485 | 7.66 | 0.006 |
| Residuals | 228 | 34.48 | 0.15 | | | 228 | 1592320 | 6984 | | |

**Table S4. Lifespan.** The effect of juvenile temperature treatment (15°C, 20°C, 25°C), strain (wild-type, *trpa-1*), and their interactions on lifespan, calculated as days since the end of the juvenile treatment.

| **Parameter** | **Loglink** | **$\chi 2$** | **d.f.** | **p** |
|---|---|---|---|---|
| Null | -1512.9 | | | |
| Treatment | -1512.6 | 0.4893 | 2 | 0.783 |
| Strain | -1512.6 | 0.1054 | 1 | 0.746 |
| Treatment × Strain | -1512.2 | 0.7273 | 2 | 0.695 |

**Table S5. Adult lifespan excluding 20°C.** The effect of juvenile temperature treatment (15°C, 25°C), strain (wild-type, *trpa-1*), and their interactions on lifespan, calculated as days since the end of the juvenile treatment.

| **Parameter** | **Loglink** | **$\chi 2$** | **d.f.** | **p** |
|---|---|---|---|---|
| Null | - 885.56 | | | |
| Treatment | - 885.52 | 0.0919 | 1 | 0.762 |
| Strain | - 885.45 | 0.1241 | 1 | 0.725 |
| Treatment × Strain | - 885.06 | 0.7762 | 1 | 0.378 |

**Table S6. Individual fitness with the differential length of the juvenile treatment taken into account.** The effect of juvenile temperature treatment (15°C, 20°C, 25°C), strain (wild-type, *trpa-1*), and their interactions on individual fitness.

| **Parameter** | **d.f.** | **SS** | **MS** | **F** | **p** |
|---|---|---|---|---|---|
| Treatment | 2 | 349.08 | 174.54 | 1218.14 | <0.001 |
| Strain | 1 | 2.34 | 2.34 | 16.35 | <0.001 |
| Treatment × Strain | 2 | 0.93 | 0.47 | 3.25 | 0.040 |
| Residuals | 339 | 48.57 | 0.14 | | |

**Table S7. Lifespan with the differential length of the juvenile treatment taken into account.** The effect of juvenile temperature treatment (15°C, 20°C, 25°C), strain (wild-type, *trpa-1*), and their interactions on lifespan.

| **Parameter** | **Loglink** | **$\chi 2$** | **d.f.** | **p** |
|---|---|---|---|---|
| Null | -1513.3 | | | |
| Treatment | -1508.8 | 9.0646 | 2 | 0.011 |
| Strain | -1508.7 | 0.1610 | 1 | 0.688 |
| Treatment × Strain | -1508.3 | 0.7393 | 2 | 0.691 |

**Analyses without removing the 5 outlier worms**

In the following tables, we have included the five outlier worms that are marked in bold in supplementary figure 1. Inclusion of these outliers does not alter the results.

**Table S8. Size with the five outliers included.** The effect of juvenile temperature treatment (15°C, 20°C and 25°C), strain (wild-type, *trpa-1*), Day, $Day^2$ and their interactions on the daily size (measured as area).

| **Parameter** | **χ2** | **d.f.** | **p** |
|---|---|---|---|
| (Intercept) | 443.00 | 1 | <0.001 |
| Treatment | 46.95 | 2 | <0.001 |
| Strain | 6.83 | 1 | 0.009 |
| Day | 2627.93 | 1 | <0.001 |
| $Day^2$ | 1855.59 | 1 | <0.001 |
| Treatment × Strain | 31.59 | 2 | <0.001 |
| Treatment × Day | 2.22 | 2 | 0.330 |
| Strain × Day | 0.38 | 1 | 0.539 |
| Treatment × $Day^2$ | 3.62 | 2 | 0.164 |
| Strain × $Day^2$ | 8.89 | 1 | 0.003 |

**Table S9. Growth rate with the five outliers included.** The effect of juvenile temperature treatment (15°C, 20°C and 25°C), strain (wild-type, *trpa-1*), Day, $Day^2$ and their interactions on the daily growth rate, with Size included as a covariate to control for size-specific growth rate.

| **Parameter** | **$\chi 2$** | **d.f.** | **p** |
|---|---|---|---|
| (Intercept) | 6784.71 | 1 | <0.001 |
| Treatment | 14.18 | 2 | <0.001 |
| Strain | 8.63 | 1 | 0.003 |
| Day | 14.64 | 1 | <0.001 |
| $Day^2$ | 0.49 | 1 | 0.482 |
| Size | 42.78 | 1 | <0.001 |
| Treatment × Strain | 13.61 | 2 | 0.001 |
| Treatment × Day | 15.67 | 2 | <0.001 |
| Strain × Day | 2.44 | 1 | 0.118 |
| Treatment × $Day^2$ | 13.76 | 2 | 0.001 |
| Strain × $Day^2$ | 2.47 | 1 | 0.116 |
| Treatment × Strain × Day | 11.44 | 2 | 0.003 |
| Treatment × Strain × $Day^2$ | 10.66 | 2 | 0.005 |

**Table S10. Development-independent reproductive rate and lifetime reproductive success, with the five outliers included**. The effect of juvenile temperature treatment (15°C, 20°C and 25°C), strain (wild-type, *trpa-1*), and their interactions on individual fitness and lifetime reproductive success.

| **Individual fitness** | | | | | | **Lifetime reproductive success** | | | | |
|---|---|---|---|---|---|---|---|---|---|---|
| **Parameter** | **d.f.** | **SS** | **MS** | **F** | **p** | **d.f.** | **SS** | **MS** | **F** | **p** |
| Treatment | 2 | 0.62 | 0.31 | 2.24 | 0.108 | 2 | 94630 | 47315 | 8.35 | <0.001 |
| Strain | 1 | 6.33 | 6.33 | 45.83 | <0.001 | 1 | 162849 | 162849 | 28.74 | <0.001 |
| Treatment × Strain | 2 | 2.38 | 1.19 | 8.62 | <0.001 | 2 | 103528 | 51764 | 9.14 | <0.001 |
| Residuals | 348 | 48.04 | 0.14 | | | 348 | 1971930 | 5666 | | |

**Table S11. Adult lifespan with the five outliers included.** The effect of juvenile temperature treatment (15°C, 20°C, 25°C), strain (wild-type, *trpa-1*), and their interactions on lifespan, calculated as days since the end of the juvenile treatment.

| **Parameter** | **Loglink** | **$\chi 2$** | **d.f.** | **p** |
|---|---|---|---|---|
| Null | -1513.0 | | | |
| Treatment | -1512.8 | 0.4893 | 2 | 0.783 |
| Strain | -1512.7 | 0.1000 | 1 | 0.752 |
| Treatment × Strain | -1512.3 | 0.7461 | 2 | 0.689 |
